# Bacterial chaperone protein Hfq facilitates the annealing of sponge RNAs to small regulatory RNAs

**DOI:** 10.1101/2021.04.25.441314

**Authors:** Ewelina M. Małecka, Daria Sobańska, Mikołaj Olejniczak

**Affiliations:** Institute of Molecular Biology and Biotechnology, Faculty of Biology, Adam Mickiewicz University, 61-614 Poznań, Poland; T.C. Jenkins Department of Biophysics, Johns Hopkins University, Baltimore, MD 21218, USA; Institute of Bioorganic Chemistry, Polish Academy of Sciences, Noskowskiego 12/14, 61-704 Poznań, Poland

**Keywords:** sRNA, small non-coding RNA, RNA chaperone, Hfq, RNA sponge, anti-sRNA

## Abstract

Bacterial small RNAs (sRNAs) in association with the chaperone protein Hfq regulate the expression of many target mRNAs. Since sRNAs’ action is crucial to engender a response to changing environmental conditions, their activity needs to be regulated. One such mechanism occurs at posttranscriptional level and involves sponge RNAs (or anti-sRNAs) which sequester sRNAs affecting their regulatory output. Both types of RNAs were identified on Hfq, but it is not known how Hfq interacts with RNA sponges and stimulates their base-pairing with sRNAs. Here, we used biochemical methods to demonstrate that anti-sRNAs resemble sRNAs by their structure and their modes of Hfq binding. Hfq facilitates sponge RNA annealing to sRNA, and each surface of the protein plays a particular role in the process. Moreover, we found that the efficiency of sponge RNA interactions with sRNAs can be improved, therefore, we propose that natural RNA sponges might not sequester sRNAs optimally.

## INTRODUCTION

Small RNAs (sRNAs) are important posttranscriptional regulators of gene expression in bacteria [1]. Most sRNAs target more than one mRNA by short and often imperfect base-pairing interactions mediated by RNA chaperone protein Hfq [2–5]. sRNA binding leads to translation activation, inhibition, and/or affects mRNA degradation [6]. This process regulates stress responses [7], virulence [8], or quorum sensing [9].

Since regulatory RNAs have a huge impact on cell metabolism, their activity needs to be tightly regulated. They are usually expressed only under specific conditions, for example transcription factors GcvA and GcvR induce GcvB synthesis in response to endogenous glycine [10]. Regulatory RNAs can be also controlled at post-transcriptional level. In eukaryotes, the activity of microRNAs (miRNAs) can be tuned after transcription by competing endogenous RNAs (ceRNAs, “sponge RNAs”). Such sponge RNAs have sequence motifs complementary to miRNA seed sequence; hence, they mimic target mRNAs [11]. The base-pairing between sponge RNA and miRNA leads to the sequestration of the latter and inhibits its activity changing regulatory output for target mRNAs. It has been recently discovered that bacterial sRNAs can be regulated in a similar way [12].

Bacterial RNA sponges (or anti-sRNAs) can be produced in multiple ways. Some of them are encoded as intergenic regions in polycistronic transcripts, e. g. SroC targeting GcvB sRNA is excised from prematurely terminated *gltIJKL* mRNA [13], while *chbBC* depleting ChiX comes from *chbB*–*chbC* spacer [14]. On the other hand, ChiZ, which also sequesters ChiX, is derived from 5’-UTR of *chiP* mRNA [15]. Other sponges are processed from the internal or external transcribed spacers (ITS or ETS, respectively) of tRNA operons. The 3′ETS of the *glyW-cysT-leuZ* mRNA (3′ETS^leuZ^) is produced via RNase E-mediated processing, and it base-pairs with two sRNAs, RyhB and RybB, providing physiological link between iron metabolism (regulated by RyhB) and membrane stress (controlled by RybB) [16]. Lastly, sponges can derive from independently transcribed genes such as AgvB and AsxR identified within bacteriophage-derived regions of the enterohemorrhagic *Escherichia coli* [17]. AgvB base-pairs with GcvB, which is one of the most globally acting sRNAs. Its targets include transcripts encoding transporters for amino acids and short peptides, proteins involved in amino acid biosynthesis and transcription factors [18, 19]. The effect of anti-sRNAs likely depends on several factors, including relative transcription rates of sponge RNAs and sRNAs, and the kinetics and binding affinities of interacting RNAs [20].

In cells, numerous sRNAs are bound by the Hfq chaperone, which stabilizes them against turnover and facilitates their annealing to mRNAs [3]. For annealing, Hfq binds both sRNAs and mRNAs by recognizing specific sequence motifs in each RNA. The distal side of Hfq binds A-rich motifs, including ARN repeats [21–23], the proximal side binds U-rich motifs [24–27], while the rim recognizes AU‑rich sequences [28, 29]. Hence, sRNAs containing Rho-independent terminators and long 3′ terminal oligo(U) tails are bound by the proximal side of Hfq [30]. Moreover, Hfq contacts sRNA “body”, but since they are highly diverse in sequence, it can occur either via rim (class I sRNAs) or distal face (class II sRNAs) of Hfq [31, 32]. Finally, mRNAs bound by sRNAs can either bind to Hfq via the distal face, when they contain A-rich motifs, or via the rim when they contain internal AU-rich sequences [2, 31]. Hfq can also exert regulatory function by binding only to mRNAs using the distal face [33–35]. Interestingly, anti-sRNAs resemble some sRNA features, e. g. they usually have 3′ oligo(U) tails, even when processed from intergenic regions. This suggests that they could interact with Hfq. Indeed, AgvB was recovered by UV crosslinking with Hfq [17], while 3’ETS^LeuZ^ was present in several RNA chimeras identified on Hfq and on another RNA-binding protein ProQ using RIL-seq [36]. The fact that SroC-mediated depletion of GcvB required the Hfq protein [13], and that RbsZ sponge downregulated RybB sRNA in the presence of Hfq [36] suggest that beyond anti-sRNA binding Hfq could also be involved in promoting their pairing to their sRNA targets. This raises a question of how Hfq mediates their interactions.

Here we present biochemical evidence that anti-sRNAs AgvB and 3′ETS^leuZ^ and their target sRNAs interact with Hfq similarly to Class I and Class II sRNAs and their mRNA targets. We also show that annealing between RNA sponges and their target sRNAs is facilitated by several Hfq surfaces, and each of them plays a particular role in this process. Moreover, we prove that the efficiency of AgvB annealing to GcvB can be enhanced indicating that natural anti-sRNA is not optimized to provide tight regulation of sRNA.

## RESULTS

### Anti-sRNAs and their sRNA targets are bound differently by Hfq

To understand the role of Hfq in mediating annealing between sRNAs and anti-sRNAs, it is a prerequisite to establish how each RNA binds this protein in the absence of the complementary partner. To explore how Hfq interacts with anti-sRNAs, we measured the affinity of AgvB and 3′ETS^leuZ^ to wild-type (wt) Hfq and its mutants with substitutions deactivating specific RNA binding surfaces: proximal face (K56A), distal face (Y25D), and the rim of the Hfq ring (R16A) (Table 2). Both anti-sRNAs showed tight, subnanomolar affinities to wild-type Hfq (Table 1) similarly to previously studied sRNAs [37, 38].

**Table 1.** Equilibrium binding of RNAs to wt Hfq and its variants with distal mutation (Y25D), rim mutation (R16A) and proximal mutation (K56A). n.m., not measured.

|  | WT | Y25D | R16A | K56A |
| --- | --- | --- | --- | --- |
| | $K_d$ (nM) | | | |
|  | (Hill coefficient) |  |  |  |
| AgvB | $0.38 \pm 0.06$<br>($1.4 \pm 0.05$) | $0.13 \pm 0.003$<br>($1 \pm 0.2$) | $2.7 \pm 0.12$<br>($0.62 \pm 0.006$) | $27.2 \pm 2.4$<br>($1.5 \pm 0.12$) |
| AgvB-U<sub>9</sub> | $0.18 \pm 0.02$<br>($1.5 \pm 0.1$) | $0.12 \pm 0.016$<br>($2 \pm 0.11$) | $0.19 \pm 0.02$<br>($1.2 \pm 0.12$) | n.m. |
| AgvB-U<sub>0</sub> | $1.75 \pm 0.6$<br>($0.7 \pm 0.05$) | $21 \pm 8$<br>($0.62 \pm 0.09$) | $3.9 \pm 0.76$<br>($0.75 \pm 0.08$) | n.m. |
| 3'ETS<sup>leuZ</sup> | $0.22 \pm 0.008$<br>($1.3 \pm 0.05$) | $0.34 \pm 0.18$<br>($0.6 \pm 0.12$) | $0.17 \pm 0.08$<br>($0.9 \pm 0.2$) | $6.8 \pm 2.9$<br>($1 \pm 0.05$) |
| 3'ETS<sup>leuZ</sup>-U<sub>4</sub> | $0.26 \pm 0.07$<br>$1.5 \pm 0.06$ | $7.3 \pm 1.3$<br>$1 \pm 0.15$ | $0.26 \pm 0.01$<br>$1.1 \pm 0.02$ | $6.3 \pm 1.8$<br>$1 \pm 0.05$ |
| RybB | $0.22 \pm 0.08$<br>($2.2 \pm 0.4$) | $0.07 \pm 0.01$<br>($1.8 \pm 0.03$) | $2.9 \pm 0.11$<br>($0.68 \pm 0.02$) | $10.2 \pm 2.5$<br>($1.9 \pm 0.04$) |
| GcvB | $0.27 \pm 0.03$<br>($1.5 \pm 0.13$) | $0.08 \pm 0.003$<br>($1.3 \pm 0.09$) | $21.8 \pm 8.5$<br>($0.5 \pm 0.03$) | $10.6 \pm 3.3$<br>($1 \pm 0.22$) |

**Table 2.**
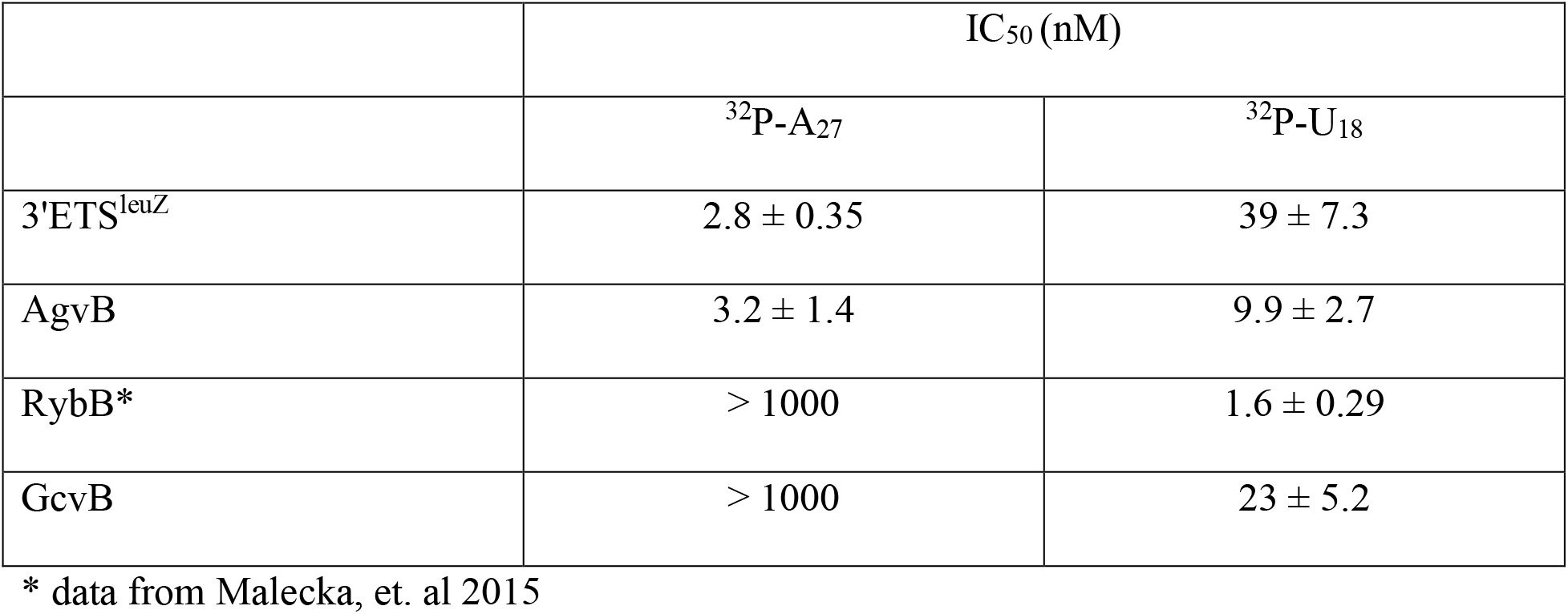
The average IC50 values for the competition of unlabeled RNAs against ^32^P-A27 and ^32^P-U18.

|  | IC<sub>50</sub> (nM) |  |
| --- | --- | --- |
|  | <sup>32</sup>P-A<sub>27</sub> | <sup>32</sup>P-U<sub>18</sub> |
| 3'ETS<sup>leuZ</sup> | 2.8 ± 0.35 | 39 ± 7.3 |
| AgvB | 3.2 ± 1.4 | 9.9 ± 2.7 |
| RybB* | > 1000 | 1.6 ± 0.29 |
| GcvB | > 1000 | 23 ± 5.2 |

\* data from Malecka, et. al 2015

Despite the fact that both AgvB and 3′ETS^leuZ^ contain single-stranded ARN repeats (Fig. 1 A, B), we did not observe significant difference in affinity to the distal face Hfq mutant as compared to wt Hfq (Fig. 1C, D, Table 1). We wondered, if the potential effect of this mutation is masked by the compensating interactions with the proximal face of Hfq. Similar masking effect was observed before, when the interactions of DsrA and RybB with the Hfq rim were undetected, until the binding to the proximal site was blocked [38]. To test if this is also the case for anti-sRNAs, we compared the binding of anti-sRNA mutants with truncated 3’-terminal oligo(U) tails: AgvB-U_0_ and 3′ETS^leuZ^-U_4_. The binding of these shorter anti-sRNAs to wt protein was only slightly affected (Fig. 1C, D, black vs. gray plots) indicating that the contacts via oligo(U) tails are not essential for binding to this protein. Consistently, the binding of AgvB-U_0_ and 3′ETS^leuZ^-U_4_ to the distal face Hfq mutant was markedly impaired compared to wt protein (Fig. 1C, D, dark red vs. light red plots). This supports the conclusion that both anti-sRNAs bind the distal face of Hfq.

**Figure 1.**
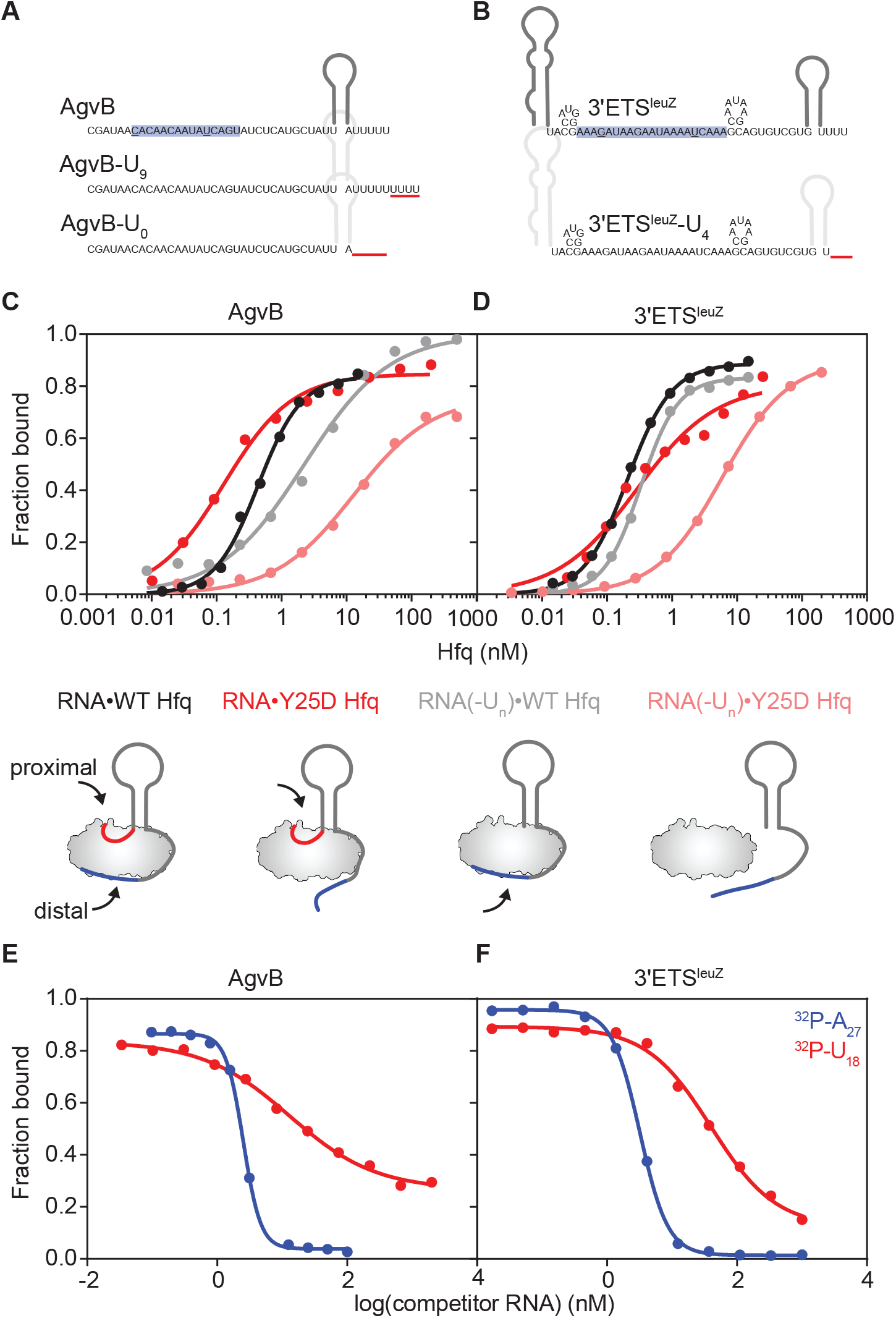
Anti-sRNAs contact multiple Hfq surfaces. (A-B) Schematic representation of (A) AgvB and (B) 3′ETS^leuZ^ structures and their mutants used in the study. Sequences added or removed are underlined in red in RNA derivatives. The longest stretch of (ARN) motifs allowing two mismatches is marked in blue, while mismatches are underlined in black. (C-D) The binding of (E) ^32^P-AgvB, ^32^P-AgvB-U_0_ or (F) ^32^P-3′ETS^leuZ^, ^32^P-3′ETS^leuZ^-U_4_ to wt Hfq (black or gray) or distal Y25D mutant (red or light red). The average *K*_d_ values can be found in Table 2. Cartoons represent schematics of RNA-Hfq contacts for each binding reaction. (E-F) The dissociation of ^32^P-A_27_ (blue) or ^32^P-U_18_ (red) from Hfq upon increasing concentrations of (C) AgvB or (D) 3′ETS^leuZ^. The average IC_50_ values can be found in Table 1.

The binding of both anti-sRNAs to the proximal face mutant was significantly weaker than to wt Hfq (Table 1), which might suggest that the interactions with distal surface are not sufficient to compensate for the lack of binding to proximal surface. Alternatively, it could indicate that RNA binding to proximal surface guides conformational changes that enable interactions with distal site in a cooperative manner. Our data indicate that both anti-sRNAs bind the distal as well as the proximal face of Hfq. However, we cannot conclude if the binding to both faces of Hfq happens simultaneously or reflects two independent binding events.

Rim mutation did not influence 3′ETS^leuZ^ binding to Hfq, but slightly decreased AgvB affinity to the protein (Table 1). To check, if the AgvB interactions with Hfq can be changed, we tested another derivative where 3′-oligo(U) tail was extended by four uridine residues (hereafter AgvB-U_9_). The binding of this RNA derivative was not dependent on the Hfq rim, which shows that stronger interactions with proximal site can alleviate the dependence on other binding surfaces.

To understand how anti-sRNAs interact with their sRNA targets on Hfq, it is also important to learn how these sRNAs bind to Hfq. Previous studies established that the target of 3′ETS^leuZ^, which is RybB sRNA, binds Hfq *via* interactions with proximal and rim surfaces [38]. To explore how Hfq binds to the target of AgvB, which is GcvB sRNA, the binding of GcvB to wt Hfq and its mutants was compared. The *K*_d_ value of GcvB binding to the wt protein was in the subnanomolar range (Table 1). Similarly to RybB, the binding of GcvB was not affected by the mutation in the distal face of Hfq, while the mutation in the proximal face was detrimental for its binding. The binding of GcvB to Hfq was more strongly affected by the mutation in the rim of Hfq than the binding of RybB (Table 1). The binding of GcvB to R16A Hfq mutant was 30-fold weaker than to wt Hfq, while the binding of RybB to R16A Hfq was 10-fold weaker than to wt Hfq. Overall, the binding of GcvB to R16A mutant was 7-fold weaker than the binding of RybB to this Hfq mutant (Table 1). These results show that besides the contacts with the proximal face of Hfq, the interactions with the rim of Hfq also play important roles for both RybB and GcvB binding to Hfq; however, they have much bigger contribution to the binding of GcvB.

Overall, these data indicate that anti-sRNAs 3′ETS^leuZ^ and AgvB interact with Hfq through contacts with at least two protein surfaces, proximal and distal. On the other hand, their target sRNAs RybB and GcvB, respectively, bind Hfq using contacts with the proximal face and the rim of Hfq.

### Both anti-sRNAs displace oligoadenylates more efficiently than oligouridylates from the complex with Hfq

To further explore how AgvB and 3′ETS^leuZ^ anti-sRNAs and their sRNA targets bind to Hfq, we analyzed their ability to displace oligonucleotides bound specifically to the Hfq distal face (oligoadenylate A_27_) or proximal face (oligouridylate U_18_) (Table 2) (Fig. 1E, F). This approach was used before to show that ChiX and MgrR sRNAs bind both the distal face and the proximal face of Hfq [38, 39], while RybB sRNA binds only to the proximal face of Hfq, in agreement with previous studies [29, 31]. Our data showed that both AgvB and 3′ETS^leuZ^ easily induce the dissociation of A_27_-Hfq complexes, with low IC_50_ values (3.2 ± 1.4 nM and 2.8 ± 0.35 nM, respectively, Fig. 1E, F). Such values are similar as previously observed for the dissociation of A_27_-Hfq complexes induced by class II sRNAs ChiX and MgrR [38, 39]. Both anti-sRNAs could displace oligouridylate U_18_ from Hfq, but at a much higher concentrations than A_27_ with IC_50_ value of 9.9 ± 2.7 nM for AgvB and 39 ± 7.3 nM for 3′ETS^leuZ^ (Fig. 1E, F), while only 4 nM of ChiX was sufficient to displace 50% of the oligouridylate from the complex with Hfq (Malecka, Strozecka et al. 2015). This result might be explained by shorter single-stranded length of 3′ oligo(U) tails present in both anti-sRNAs, which is five nucleotides long for AgvB and three for 3′ETS^leuZ^, as compared to seven-nucleotide long single-stranded 3’-tail of ChiX sRNA.

It has been previously reported that RybB specifically binds to the proximal face of Hfq, which is shown by the fact that it exclusively displaces U_18_, but not A_27_ from Hfq [38]. To compare this behavior with that of GcvB, we analyzed whether GcvB can displace U_18_ and A_27_ from Hfq. The data showed that GcvB was not able to outcompete oligoadenylate from Hfq even at high concentrations indicating that it does not interact with distal surface (Table 1). Surprisingly, its ability to displace U_18_ from Hfq was also much lower than for RybB. The previously reported IC_50_ value for RybB was 1.6 ± 0.29 nM, while the value for GcvB determined here was 23 ± 5.2 nM (Table 1).

Overall, these data showed that despite binding to both the distal and proximal face of Hfq (Table 1), anti-sRNAs 3′ETS^leuZ^ and AgvB are much more efficient at displacing A_27_ from the distal face than U_18_ from the proximal face (Table 2), which suggests that the distal face is their main site of binding to Hfq. It contrasts with the behavior of sRNAs RybB and GcvB targeted by tested sponges.

### AgvB and 3′ETS^leuZ^ contain long ARN sequence repeats located in single-stranded regions

To determine the secondary structures of anti-sRNAs AgvB and 3′ETS^leuZ^ they were probed using structure-specific RNases (Fig. 2). The positions of degradation induced by RNase T2 and Nuclease S1 were used as constraints to define single-stranded regions. Moreover, RNase T1 was used, which specifically degrades single-stranded RNA at G residues. To utilize RNase T1 to distinguish between single-stranded and double-stranded regions, we performed experiments at native (degradation at accessible Gs) and denaturing conditions (degradation at all Gs). Cleavage patterns indicated specific nucleotide positions that were used as constraints in structure predictions using *RNAstructure* software [40].

**Figure 2.**
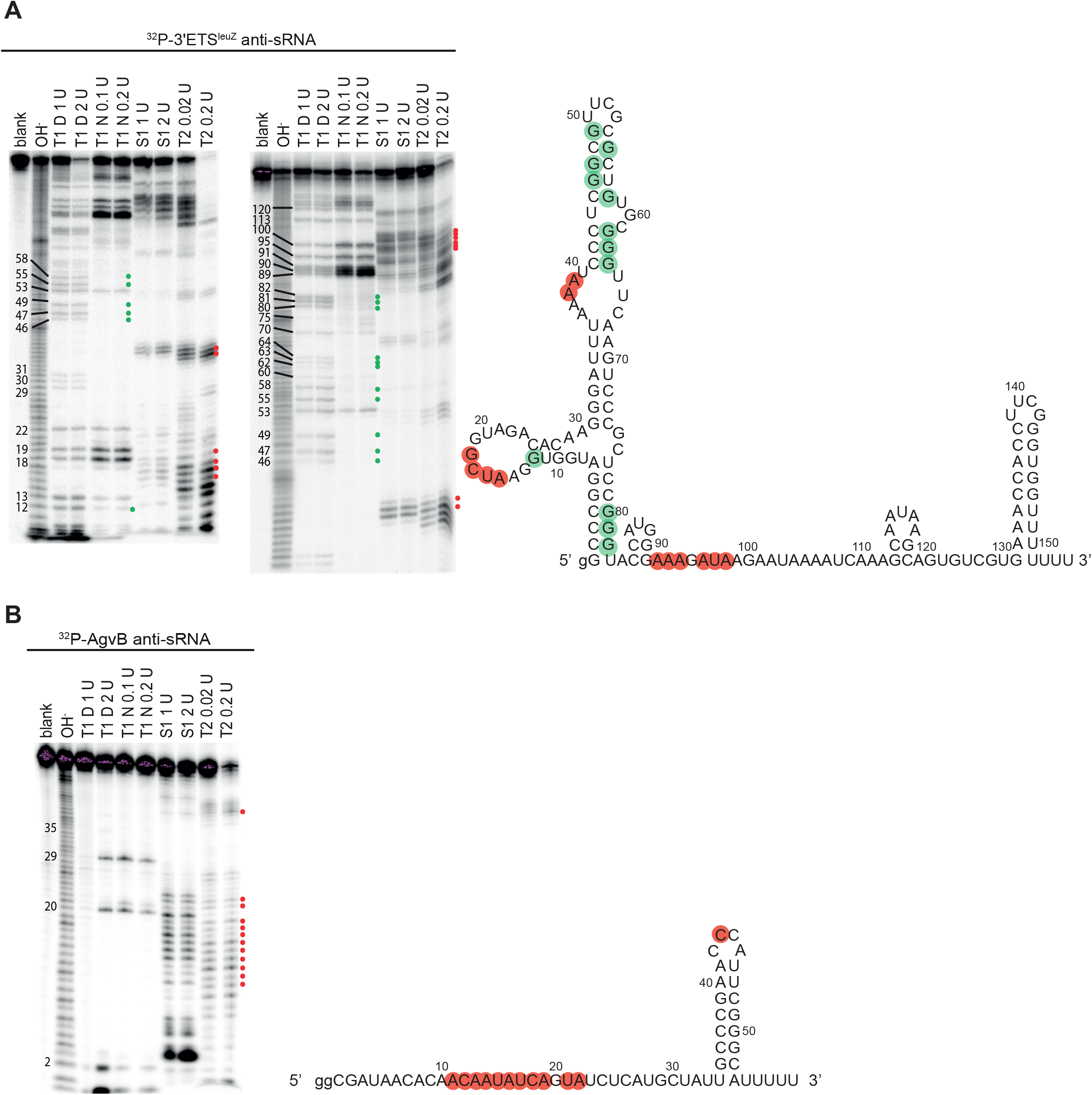
Secondary structures of 3ʹETS^leuZ^ and AgvB. (A) Left: ^32^P-labeled 3ʹETS^leuZ^ structure probing with RNases indicated above the lanes. T1 D and T1 N denote probing with RNase T1 in denaturing or native conditions, respectively. The numbers on the left indicate positions of guanosine residues. Blank points out untreated control and OH-denotes formamide ladder. Right: 3ʹETS^leuZ^ secondary structure predicted by RNAstructure software based on the data from structure probing experiments showed on the left. Guanosine residues not cleaved by RNase T1 in native conditions were constrained as double-stranded (green circles). Residues cleaved with RNase T2 or Nuclease S1 were constrained as single-stranded (red circles). (B) Left: ^32^P-AgvB structure probing. Right: AgvB secondary structure predicted by RNAstructure software based on the data from structure probing experiments showed on the left. Figure description as in (A).

Analysis of anti-sRNAs sequences revealed that both contain repeated ARN motifs (adenine, purine, any nucleotide), which are known to direct either mRNAs or Class II sRNAs to the distal face of Hfq. Allowing some flexibility in ARN motif (max. 2 mismatches), we identified the longest stretch of ARN repeats to be 5 for AgvB (Fig. 1A) and 7 for 3′ETS^leuZ^ (Fig. 1B).

The probing data for 3′ETS^leuZ^ anti-sRNA showed the formation of two stable stem-loops in 5ʹ- and 3ʹ-terminal regions of the RNA (Fig. 2A). Our results showed that nucleotide residues located at positions 92-94 and 96-98 in the region including seven ARN repeats, are susceptible to cleavage by RNase T2 and Nuclease S1. This suggests that these repeated ARN motifs are located in a single-stranded region, which makes them potentially available for interaction with Hfq’s distal face. Additionally, the sequence of the RybB binding site (positions 107 to 127) includes a small stem-loop structure with two G-C pairs. The cleavage pattern of the 3ʹ-oligoU tail was not well resolved, although the three terminal uridines were predicted to be single-stranded and, hence, potentially accessible for binding by the proximal face of Hfq.

*In vitro* structure probing of AgvB RNA revealed that it forms a stable stem-loop at its 3ʹ-terminus (Fig. 2B). However, its 5ʹ-terminus and the central part with A-rich region are single-stranded, and therefore, accessible for Hfq binding. Moreover, the 3ʹ-terminal U-rich region was predicted to be single-stranded, and available for interaction with the proximal site of Hfq. Nucleotides involved in pairing with GcvB (positions 6 to 23) were also located in the unstructured region, potentially allowing efficient RNA-RNA duplex formation.

As opposed to both anti-sRNAs, their sRNA targets only contain 3’-terminal oligoU tails, and do not have internal ARN sequence repeats. The previously analyzed secondary structure of *S. enterica* RybB shows the presence of single-stranded 3’ terminal oligoU tail, and unstructured 5’ terminal and central region, which contains an AU-rich sequence [28, 29]. When the secondary structure of GcvB was analyzed using RNases, the data showed that GcvB contains mostly single-stranded, seven-uridine tail, while its central and 5’ terminal regions are highly structured (Fig. S1). As the central structured regions contain U-rich sequences, especially in loop and bulge regions, it suggests that these regions could be restructured by Hfq, as shown for similar motifs before [41], and hence could be available for Hfq to bind.

Overall, our probing data indicated that AgvB and 3ʹETS^leuZ^ anti-sRNAs contain both repeated ARN sequence motifs located in their central regions, and Rho-independent transcription terminator hairpins followed by oligoU tails. The presence of these two kinds of sequence motifs makes them similar to Class II sRNAs [31], but it also suggests that they could bind to Hfq either similarly as an sRNA or similarly as an mRNA.

### The Hfq-dependent association of 3′ETS^leuZ^ to RybB is most strongly disrupted by the mutations in the distal face and the rim of Hfq

To elucidate the role of Hfq in the annealing of sRNAs to anti-sRNAs, the kinetics of association was monitored by electrophoretic mobility shift assay. The annealing of ^32^P-RybB to 3′ETS^leuZ^ (Fig. 3A) was accelerated in the presence of wt Hfq, similarly to previous observations for sRNA-mRNA pairs [42, 43]. The rate of Hfq-mediated annealing was increased 90-fold (*k*_obs_ 0.022 ± 0.04 min^−1^ without Hfq vs. 1.98 ± 0.31 min^−1^ in the presence of 5 nM Hfq, Fig. 3B, C, Table 3).

**Figure 3.**
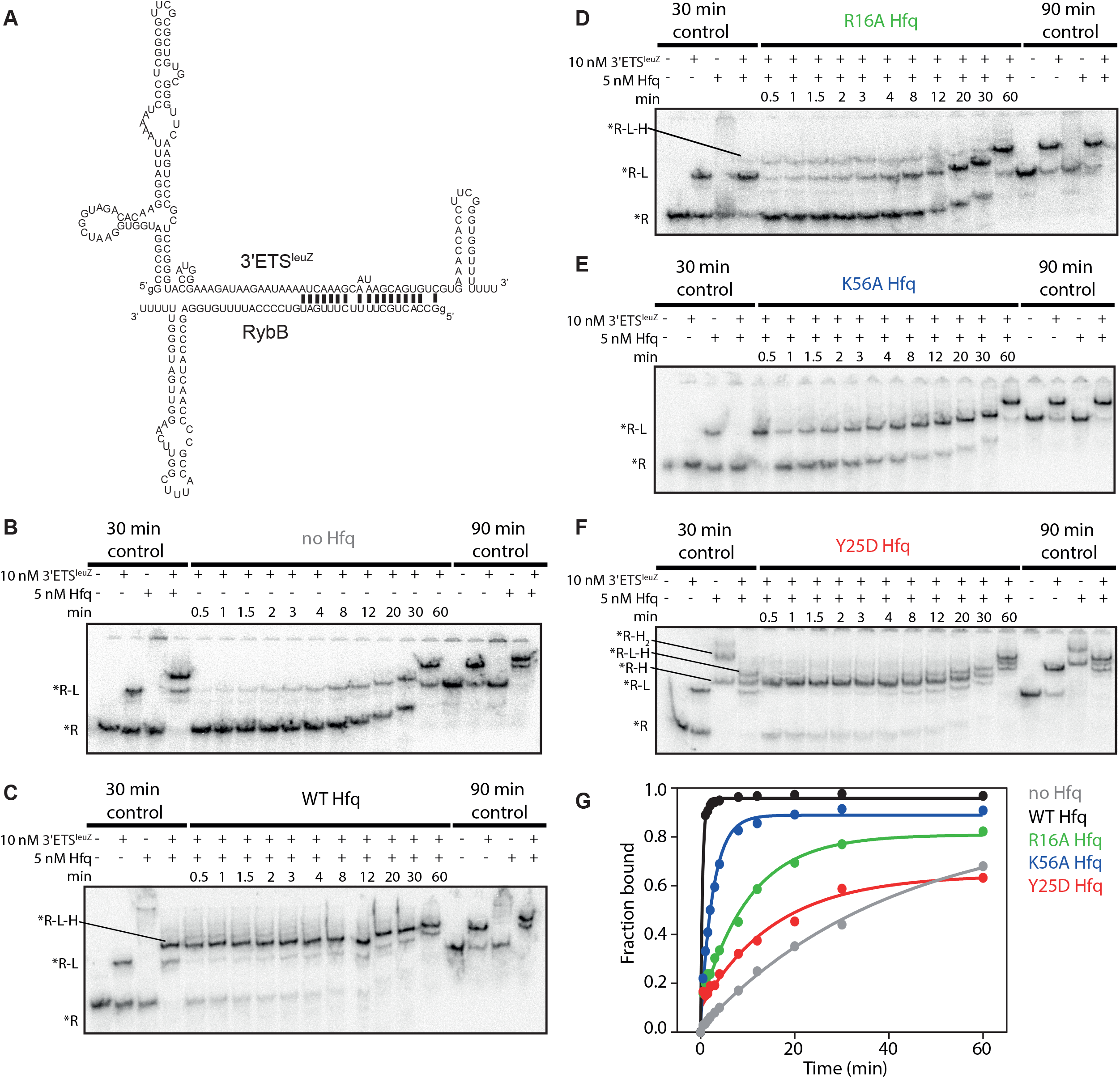
RybB-3′ETS^leuZ^ annealing is strongly inhibited by Hfq distal mutation. (A) Base-pairing between 3′ETS^leuZ^ and RybB. (B-F) The kinetics of ^32^P-RybB annealing to 3′ETS^leuZ^ in the absence or presence of 5 nM Hfq or its mutants as indicated. Free ^32^P-RybB is marked as R, 3′ETS^leuZ^ as L, and Hfq as H. Combinations of those letters refer to their complexes. (C) The complex of ^32^P-RybB with wt Hfq was not well resolved. To double check if the composition of complexes is assigned properly, a control reaction was performed, and the gel was resolved in TBE buffer. It confirmed that the slowest migrating complex formed in the presence of both RNAs and Hfq is a ternary complex. For details see Fig. S2. (G) The data from B-F were plotted versus time and fitted with single exponential equation. Average *k*_obs_ values can be found in Table 3.

**Table 3.** The rates of sRNA-anti-sRNA annealing in the presence of wt Hfq and its variants.

|  | *k*<sub>obs</sub> (min<sup>-1</sup>) |  |  |  |  |
| --- | --- | --- | --- | --- | --- |
|  | - Hfq | + wt Hfq | + Hfq Y25D | + Hfq R16A | + Hfq K56A |
| <sup>32</sup>P-RybB-3'ETS<sup>leuZ</sup> | 0.022 ± 0.004 | 1.98 ± 0.31 | 0.059 ± 0.013 | 0.11 ± 0.011 | 0.37 ± 0.004 |
| <sup>32</sup>P-AgvB-GcvB | 0.026 ± 0.01 | 0.55 ± 0.15 | 0.023 ±<br>0.0007 | 0.041 ± 0.02 | 0.05 ± 0.006 |

To assess the contribution of each Hfq surface to the base-pairing process, we also monitored the association in the presence of Hfq mutants. Rim mutant increased the annealing rate only 5-fold compared to the reaction without Hfq (Fig. 3D). A fraction of the ternary complex could still be detected on the gel indicating that RNAs can coexist relatively stably on Hfq. It agrees with the observation that the Hfq rim is not essential for the binding of either of the two RNAs: RybB is anchored on the proximal surface, while 3′ETS^leuZ^ on the distal surface of Hfq (Figure 1, Table 1, 2). The reason for a slow annealing rate might be the lack of positively charged arginine on the rim which has been shown to catalyze the base-pairing reaction [44]. As a result, helix nucleation might not occur efficiently, which leads to complex dissociation.

Partial, 17-fold stimulation of base-pairing was also retained in the presence of proximal face mutant (Fig. 3E). However, we did not observe the ternary complex on the gel. It might suggest that Hfq can still facilitate annealing reaction, but the stability of ternary complex is impaired and base-paired RNAs are quickly released.

Finally, we noted only 2-fold increase in annealing rate in the presence of the distal face mutant (Fog. 3F). Since this surface is important for binding 3′ETS^leuZ^, we expected that the anti-sRNA recruitment process is impaired when distal site is not available. In agreement with that, we observe a high fraction of RybB-Hfq complex on the gel, but no binary complex of Hfq with 3′ETS^leuZ^.

Overall, the results show that the interactions of RybB with the proximal face of Hfq are important to maintain the stability of the ternary complex. Additionally, 3′ETS^leuZ^ is recruited to the distal site, while the rim of Hfq stimulates the duplex formation. Hence, while RybB binds Hfq as a Class I sRNA, the binding mode of 3′ETS^leuZ^ is similar to that of an mRNA regulated by a Class I sRNA (Schu, Zhang et al. 2015).

### The Hfq-dependent annealing of AgvB to GcvB is strongly susceptible to mutations in all three RNA binding sites of Hfq

The annealing of ^32^P-AgvB to GcvB (Fig. 4A) was accelerated in the presence of Hfq (Fig. 4B, C), In the absence of Hfq, the two RNAs annealed with a rate of 0.026 ± 0.01 min^−1^, while in presence of 3 nM Hfq they interacted 20 times faster at a rate of 0.55 ± 0.15 min^−1^ (Fig. 4G, Table 3).

**Figure 4.**
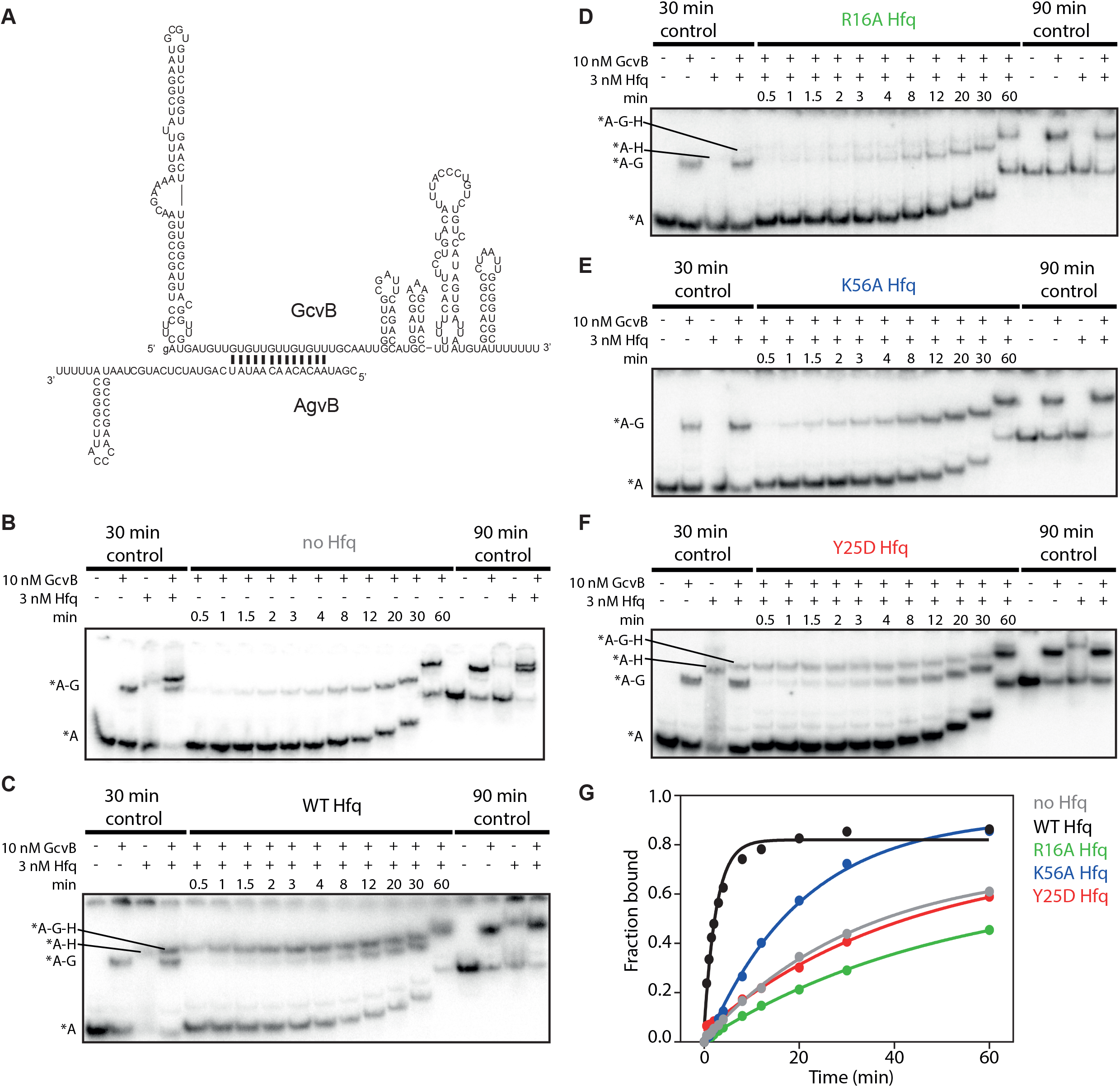
The interaction of AgvB with GcvB is strongly inhibited by Hfq mutations. (A) Base-pairing between AgvB and GcvB. (B-F) The kinetics of ^32^P-AgvB annealing to GcvB in the absence or presence of 3 nM Hfq or its mutants as indicated. Free ^32^P-AgvB is marked as A, GcvB as G, and Hfq as H. Combinations of those letters refer to their complexes. (G) The data from B-F were plotted versus time and fitted with single exponential equation. Average *k*_obs_ values can be found in Table 3.

Annealing acceleration was abolished when reaction was performed with the Hfq rim mutant (Fig. 4D). As we showed above (Table 1) this surface is important for GcvB binding to Hfq. Additionally, it could contribute to interactions with AgvB. Consistently with that, Hfq-mediated annealing is inhibited, and no ternary complex was formed.

Similarly, the proximal face mutant of Hfq showed only 2-fold increase in the association rate (Fig. 4E). This surface is bound by AgvB, which can explain the impaired annealing. Even though AgvB also contacts distal surface, these interactions were not sufficient for RNA to remain bound to Hfq and anneal to GcvB. In fact, the sequence of the distal site binding motif in AgvB overlaps with the sequence partially complementary to GcvB. It indicates that AgvB interactions with proximal surface are dominant during the engagement with GcvB, and provide complex stability. It is consistent with lack of ternary complex on the gel in the presence of proximal mutant.

Interestingly, we also observed very slow annealing in the presence of the distal face mutant (Fig. 4F). Both GcvB and AgvB efficiently bind the distal face mutant with subnanomolar *K*_d_ values (Table 1). Therefore, the low affinity to the protein does not explain slow association rate. In fact, the formation of ternary complex can be observed on the gel suggesting that some binding encounters are successful. The sequence motif in AgvB predicted to bind distal surface overlaps with region complementary to GcvB. Therefore, it is possible that the interactions between the A-rich region of AgvB and the distal face of Hfq are necessary to ensure that the seed sequence in AgvB is properly oriented to efficiently base-pair with GcvB.

Overall, these results indicate that AgvB contacts with proximal site of Hfq provide stability of AgvB/Hfq complex, while GcvB is engaged through the rim, which is similar to the mode of binding of Class II sRNAs and their mRNA targets [31]. Interactions with the distal site might play an important role in proper arrangement of AgvB seed sequence on Hfq in a way that is poised for base-pairing with GcvB.

### Strengthening AgvB interactions with Hfq proximal site accelerates its annealing with GcvB

We showed that the mode of AgvB and GcvB binding to Hfq involves the same protein surfaces. Both AgvB and GcvB displayed the ability to interact with proximal site, although the affinity was lower than for other sRNAs (Fig. 1C, D). Moreover, their binding also depended on the Hfq rim. Previous studies suggested that RNA interactions with the same binding surface would lead to displacement of one of them from the protein [38, 45]. Even though we observed the formation of ternary complex with wt Hfq, we wondered if the annealing process was efficient. Hypothetically, some encounters of incoming RNA with RNA-Hfq complex could be unsuccessful because the preferred binding site is already occupied by the resident RNA. We wondered whether strengthening RNA interactions with one Hfq binding surface could alleviate the dependance on the other site and thus increase the efficiency of Hfq-mediated annealing. We already noticed that extending 3′ oligo(U) in AgvB, and thus strengthening its interactions with proximal surface, relieved AgvB dependency on interactions with the rim, which is the main point of contact with GcvB (Table 2). Therefore, we decided to check if the efficiency of sRNA-sponge annealing can be manipulated by changing the strength of RNAs interactions with proximal Hfq surface.

To check if kinetics of Hfq-mediated AgvB-GcvB annealing is changed when AgvB has an extended 3′ oligo(U) tail, we performed gel shift assays in the presence of 9 nM wt Hfq. At this Hfq concentration AgvB annealed to GcvB with a rate 1 ± 0.16 min^−1^ (Fig. 5A, S3A). Extending AgvB 3′ oligo(U) increased this rate to 2.5 ± 0.75 min^−1^ (Fig. S3B). It suggests that stronger binding of AgvB to proximal surface enables more efficient interactions with rim-dependent GcvB. If this is the case, we should also observe a slower association when AgvB is completely dependent on the rim or distal site which should interfere with GcvB binding. To test that, we monitored the annealing of GcvB to truncated AgvB containing no uridines at 3′end (AgvB-U_0_). As expected, the observed rate 0.58 ± 0.18 min^−1^ was slower than noted for wt AgvB (Fig. S3C).

**Figure 5.**
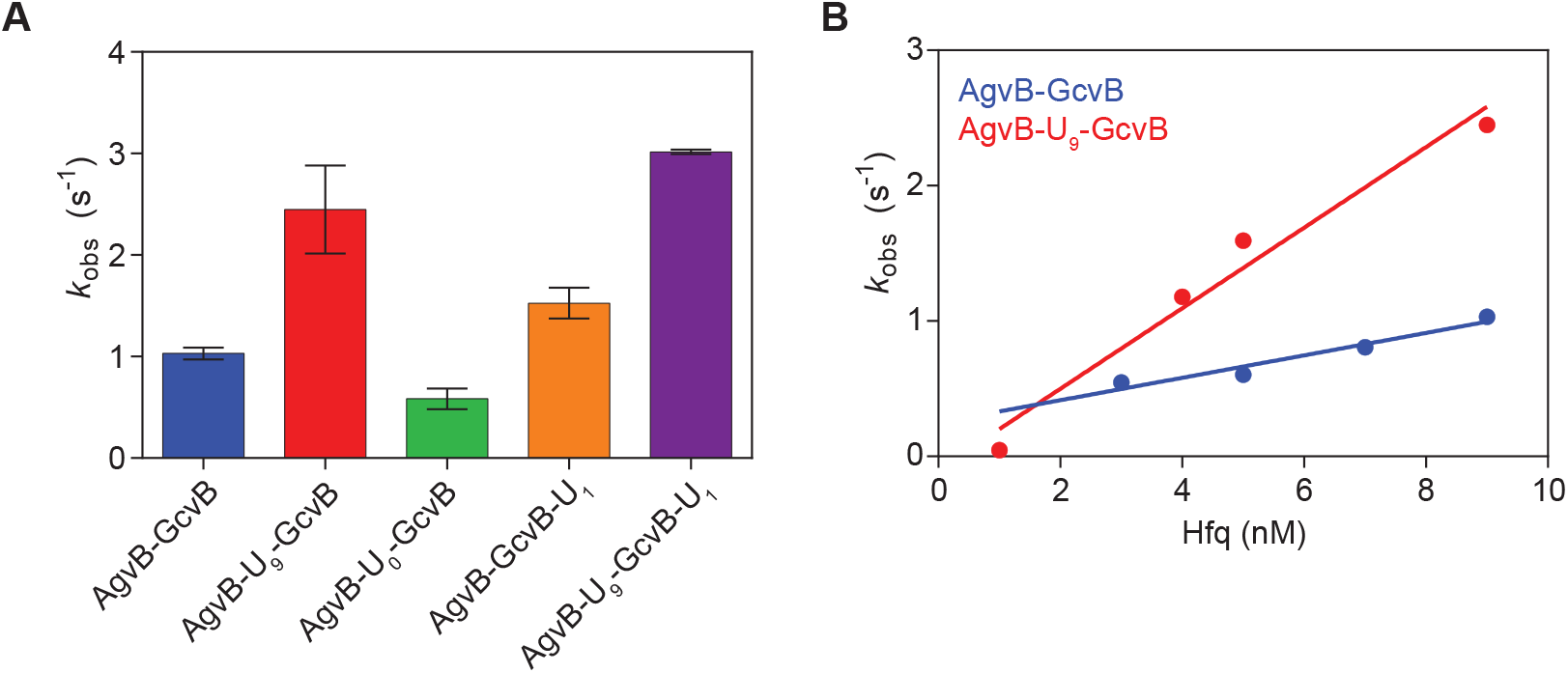
AgvB sponge does not associate with sRNA in the most efficient way. (A) Bar graph representing average *k*_obs_ values annealing between indicated RNAs in the presence of 9 nM wt Hfq. (B) Observed rates of ^32^P-AgvB-GcvB (blue) and ^32^P-AgvB-U_9_-GcvB (red) association (*k*_obs_) plotted versus Hfq concentration. The linear increase in *k*_obs_ yielded *k*_on_ value higher for AgvB with extended 3′oligo(U) (0.3 ± 0.03 nM^−1^min^−1^) than wt AgvB (0.083 ± 0.01 nM^−1^min^−1^). See also Fig. S4.

Based on the competition experiments with U_18_, GcvB was able to weakly interact with proximal surface (Table 1). Therefore, it could interfere with AgvB binding. To check that, we tested how the kinetics of annealing would change when GcvB is deprived of terminal uridines (GcvB-U_1_). We noted a slight increase in *k*_obs_ compared to wt GcvB (1.53 ± 0.2 min^−1^ vs 1 ± 0.16 min^−1^, Fig. S3D). It shows that the interactions of GcvB with proximal surface are not necessary for efficient annealing with its sponge.

Based on our findings, the most efficient annealing should occur between RNAs without any overlapping binding sites on Hfq. To check this hypothesis, we monitored the annealing of extended AgvB (AgvB-U_9_) to truncated GcvB (GcvB-U_1_). In fact, the observed rate was the fastest out of all tested RNA variants (3 ± 0.02 min^−1^, Fig. S3E).

To further confirm that our conclusions are independent of protein concentration, we performed association experiments between AgvB-GcvB and AgvB-U_9_-GcvB at a range of Hfq concentrations (Fig. 5B, Fig. S4). The linear increase in *k*_obs_ obtained for AgvB-GcvB plotted as a function of Hfq concentration yielded second order rate constant 0.083 ± 0.01 nM^−1^min^−1^. When the extended version, AgvB-U_9_ was tested, we noted higher observed association rates. Therefore, resulting *k*_on_ was also faster, 0.3 ± 0.03 nM^−1^min^−1^. Overall, the results show that strengthening the interactions of AgvB with proximal site of Hfq relieves the rim surface which now can be occupied by GcvB for efficient annealing.

## DISCUSSION

Anti-sRNAs are important regulators of sRNAs’ activity. Since sRNAs interact with chaperone protein Hfq, it is crucial to understand how the interactions with the sponge are facilitated by the RNA chaperone. Here, we dissected the mode of Hfq binding and the mechanism of Hfq-mediated annealing for two sRNA-sponge interactions: AgvB-GcvB and 3′ETS^leuZ^-RybB.

### Mode of anti-sRNAs interaction with Hfq

To determine how AgvB and 3′ETS^leuZ^ interact with Hfq, we performed competition experiments and measured equilibrium dissociation constants for binding to Hfq and its mutants. The competition with oligoadenylate revealed that both anti-sRNAs efficiently interact with distal Hfq surface. This result agrees with the previous study demonstrating that AgvB was able to displace the ARN motif-containing *dppA* mRNA from the complex with Hfq *in vitro* [17]. AgvB and 3′ETS^leuZ^ in our assays were also able to displace oligouridylate, but at higher concentrations than previously tested sRNAs, suggesting that their affinity to proximal surface is lower. This observation might be explained by the length of unpaired 3′ oligo(U) region, which in both anti-sRNAs is shorter (AgvB – 5 nt, 3′ETS^leuZ^ – 3 nt) than in typical sRNAs (e. g ChiX - 7 nt). It has been shown before that the number of terminal uridines affects the sRNAs’ ability to displace other sRNAs from Hfq [37, 38]. Moreover, systematic truncations of 3′ oligo(U) in SgrS sequence resulted in gradual decrease in sRNA stability in cells. The removal of as little as two terminal uridines (6 uridines remaining) partially impaired the regulation of *ptsG* mRNA regulation [26]. This suggests that neither of the tested anti-sRNAs utilize the interactions with the Hfq proximal site efficiently.

Even though studied sponges did not show high affinity to proximal surface, its deactivation resulted in decreased binding efficiency to the protein. Both AgvB and 3′ETS^leuZ^ are recognized by the distal surface, but these interactions were not sufficient to retain the tight binding to K56A proximal mutant. Similar effect was observed for MgrR sRNA, which also has a relatively strong ARN motif allowing for the efficient displacement of A_27_ from Hfq and thus, binding to its distal site [39]. Interestingly, the mutation Y25D in the distal Hfq surface did not impair anti-sRNAs binding. The effect of this mutation was observed only when unpaired 3′ terminal uridines were removed. It might suggest that the binding to proximal surface guides the interactions with other surfaces, and it is a universal mechanism of recognizing sRNAs or anti-sRNAs by Hfq. One exception might be GcvB sRNA, which showed unusually strong dependence of Hfq binding on the rim surface.

Taken together, our studies showed that AgvB and 3′ETS^leuZ^ anti-sRNAs occupy two surfaces of Hfq: the distal and the proximal one, which resembles the mode of class II sRNAs interaction with Hfq.

### Hfq surfaces contribute differently to sRNA-sponge base-pairing

To get an insight into the mechanism of Hfq-mediated sponge-sRNA annealing, we tested the base-pairing in the presence of Hfq mutants. The kinetics of annealing of tested RNA pairs were moderately affected by proximal K56A mutation on Hfq, which is consistent with weaker binding of sRNAs to this mutant. However, even though the protein was able to partly accelerate the base-pairing reaction, ternary complex was not detected on the gel (Fig. 3E, 4E). It indicates that the interactions with the proximal surface provide stability of ternary complex.

3′ETS^leuZ^ annealing to RybB was shut down in the presence of the distal face mutant (Fig. 3F). It has been observed before that RybB base-pairing with its target mRNA, *ompD* was impaired in the presence of the Hfq distal mutant (Wroblewska and Olejniczak 2016). Similarly to *ompD*, 3′ETS^leuZ^ contains ARN motifs which mediate interactions with this surface of Hfq explaining the role of distal side in 3′ETS^leuZ^ recruitment. The annealing of 3′ETS^leuZ^ to RybB was also slowed down by rim R16A Hfq mutation (Fig. 3D). Besides binding AU-rich sequences, e. g. in RybB, rim has been shown to catalyze the base-pairing reaction (Panja 2013). This mutation also negatively affected the base-pairing between RybB and *ompD* (Wroblewska and Olejniczak 2016), which implies that interactions between RybB and its complementary RNAs are stimulated by positively charged arginies. However, the effect of the rim mutation was less severe for RybB-*ompD* annealing than RybB-3′ETS^leuZ^. Since RybB binds both the Hfq rim and the proximal surface (Sauer 2012, Malecka 2015), sRNA interacts with proximal side when the rim is deactivated. However, 3′ETS^leuZ^ also showed the ability to bind proximal surface, although with lower affinity (Table 1). Therefore, it is possible that 3′ETS^leuZ^ might displace RybB from Hfq when sRNA binding is partly destabilized by rim mutation. It could not only cause slower effective annealing but also faster duplex release which is in fact observed on the gel (Fig. 3D). Overall, our findings indicate that RybB-3′ETS^leuZ^ annealing resembles the interactions of class I RNAs, where 3′ETS^leuZ^ is a counterpart of class I mRNA.

Since GcvB-Hfq interactions strongly depend on the rim (Table 2), it was expected that the annealing will be impaired by this mutation. In fact, the fraction of ternary complex was barely detectable on the gel in the presence of the R16A protein (Fig. 4D). In this way GcvB resembles mRNAs targeted by class II sRNAs, such as *eptB* or *chiP*, for which the rim mutant also did not stimulate annealing (Kwiatkowska et al., 2018). Like class II mRNAs, the sequence of GcvB is rich in uridines which mediate the interactions with the Hfq rim [28, 29, 46]. The distal face mutation also affected the AgvB-GcvB annealing even though both RNAs exhibited efficient binding to this mutant in equilibrium experiments. However, AgvB was found to interact with the distal side (Table 1, 2). On the other hand, the ARN motif in AgvB overlaps with the sequence base-pairing to GcvB. Indeed, it has been found before that Hfq and RNA binding motifs can overlap [17]. These interactions with Hfq might pre-expose the complementary region enabling the base-pairing, as it has been shown for RydC and rim surface [28, 29, 46]. If the distal surface is not available for AgvB binding, it is possible that its 5′ end becomes more flexible and therefore, the seed region is not pre-oriented properly. This way AgvB-GcvB annealing resembles the base-pairing mechanism of class II RNAs, where GcvB plays a role of targeted mRNA while AgvB behaves similarly as class II sRNA.

### AgvB does not sequester GcvB optimally

The analysis of kinetics of RNA-RNA association revealed that the interactions of AgvB with proximal surface are important for the stability of ternary complex during GcvB engagement. Surprisingly, AgvB interacts only weakly with this surface. In fact, strengthening AgvB interactions with proximal site increased the annealing rate (Fig. 5) indicating that natural sponge can be further optimized to provide faster base-pairing.

Even though AgvB-GcvB formed ternary complex with Hfq, we observed duplex release over time on the native gel. Interestingly, ternary complex did not dissociate when AgvB with extended 3′ oligo(U) tail was tested (Fig. S3). This indicates that strong Hfq binding motifs provide higher stability of ternary complex which might affect the downstream regulatory processes such as degradation. It has been shown that Hfq recruits degradosome (Bandyra 2012, Ikeda 2011). Interestingly, in cells AgvB antagonizes GcvB activity but does not cause sRNA destabilization (Tree 2014). Miyakoshi reported that other sponge targeting GcvB in *Salmonella*, SroC, triggers GcvB degradation (Miyakoshi 2015). SroC base-pairs with other regions of GcvB than AgvB which might be one of the factors affecting the downstream mechanism. However, SroC-mediated depletion of GcvB is Hfq dependent suggesting that the stability of ternary complex might also regulate it. SroC sponge has motifs that can provide stable binding to Hfq but it has not been studied *in vitro*. It would be interesting to investigate further what are the determinants of SroC-mediated GcvB degradation as opposed to sequestration provided by AgvB.

There are many factors that can affect the efficiency of sponges in cells, such as the transcription rate, how fast transcript can be processed in case of e. g. SroC or 3′ ETS^leuZ^, their level in the cells, and ratio to regulated sRNA. Because sRNAs are bound by Hfq, which also mediates their interaction with sponges, other crucial factors are the ability to bind the protein and to remain in a stable complex. Since AgvB-GcvB annealing could be easily optimized by improving natural AgvB sequence, we suspect that this step might be limiting for AgvB action in cells. It is possible that the role of some sponges is only to fine-tune the level of sRNAs, while other might act more efficiently in cells.

Artificial mRNA-based miRNA sponges have been used to inhibit miRNA functions in a specific way even before natural sponges have been discovered [47]. Anti-sense RNAs could be also used in bacteria to shut down sRNA regulation. Our study pinpoints the importance of Hfq binding motifs which should be considered in design of such anti-sRNAs.

## MATERIALS AND METHODS

### RNA preparation

RNAs used in this study were *in vitro* transcribed using T7 RNA polymerase [48], and purified on 10% polyacrylamide, 8M urea gel as previously described [37]. The DNA templates for transcription were obtained by Taq extension of overlapping oligonucleotides (Sigma Aldrich). RNAs were 5′-labeled with ^32^P-γ-ATP using T4 polynucleotide kinase (NEB) and purified on P-30 spin columns (Bio-Rad). Chemically synthesized oligoribonucleotides A_27_ and U_18_ were kind gifts of Prof. Ryszard Kierzek (Institute of Bioorganic Chemistry, Polish Academy of Sciences, Poznan).

### Hfq protein expression and purification

C-terminally His_6_-tagged *E. coli* wild type (wt) Hfq protein and its mutants were expressed from pET15b vector (Novagen) and purified as described [38]. Cells were grown to OD_600_~0.5, induced with 1 mM IPTG (final concentration) followed by 3-4 h at 37°C, and collected by centrifugation. Resulting pellet was resuspended in buffer A (50 mM HEPES, pH 7.5, 500 mM NH_4_Cl, 5% (w/v) glycerol) with protease inhibitor tablet (EDTA-free, Roche), and lysed by sonication. Resulting lysate was clarified by centrifugation and loaded onto HisTrap crude column charged with NiSO_4_ (GE Healthcare). Unbound proteins were washed out with buffer A supplemented with 10 mM imidazole. Proteins bound to the resin were eluted with a linear gradient of imidazole (35-600 mM) in the buffer A. To remove any residual nucleic acids, fractions containing Hfq protein were combined, concentrated, and treated with RNase A (30 μg/mL) and DNase I (5 U/mL) for 1 hour at 37°C. The solution was diluted to decrease the imidazole concentration and passed through a Ni^2+^ affinity column. Fractions after elution were concentrated and applied to a HiLoad 16/60 Superdex 200 size exclusion column (GE Healthcare) equilibrated with the storage buffer (50 mM HEPES 7.5, 250 mM NH_4_Cl, 1 mM EDTA, and 10% glycerol). After elution, the protein was concentrated and stored in aliquots at −80°C.

### Equilibrium binding assay

To determine the affinity of RNA binding to wt or mutant Hfq protein, a double filter retention assay was employed as previously described [38]. Prior to binding, RNAs were denatured in the binding buffer (50 mM Tris-HCl pH 7.5, 50 mM NaCl, 50 mM KCl, 0.1 mM EDTA) for 2 min at 90°C followed by 10 min incubation at room temperature. During refolding, MgCl_2_ was added to RNAs to obtain a 2.5 mM final concentration. To initiate binding, 0.1 nM ^32^P-labeled RNA (25 μL) was mixed with an equal volume of Hfq protein dilution in the binding buffer (in a desired range of concentrations). After 25 min incubation at room temperature, 45 μL from each reaction was passed through the filters, and immediately washed with 100 μL binding buffer. Filters were hot air dried, exposed to the phosphor screen overnight, and quantified using a phosphorimager and MultiGauge software (Fuji). The fraction of ^32^P-labeled sRNA bound to Hfq was fit to a two-state binding isotherm, ƒ_B_ = [Hfq_6_]^n^/([Hfq_6_]^n^ + *K*_d_^n^), in which n is the Hill coefficient.

### Equilibrium competition assay

Competition assays were performed as previously described [38]. In brief, RNAs were prepared as for the equilibrium binding assay. Components were mixed in a 30 μL reaction volume at final concentrations as follows: 4 nM Hfq, 0.04 nM ^32^P-labeled A_27_ or ^32^P-labeled U_18_, and a range of concentrations of unlabeled RNA. After 35 minutes of incubation, 25 μL aliquots were filtered and washed with 100 μL binding buffer. Membranes were treated as described above. The fraction of ^32^P-labeled sRNA bound to Hfq was fit using the equation 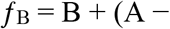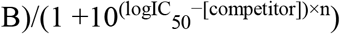, where ƒ_B_ is a fraction bound, A and B are plateaus (top and bottom, respectively) in the units of the Y axis, n is the Hill coefficient.

### Kinetics of anti-sRNA-sRNA annealing

The association kinetics of anti-sRNAs to sRNAs was monitored by native electrophoretic mobility gel shift assay as described before [43]. RNAs were prepared as for equilibrium binding assay. Binding reactions were carried out at 25°C in 120 μL volume. 1 nM ^32^P-labeled RNAs were incubated with 10 nM of unlabeled RNA in the absence or presence of indicated concentration of wt Hfq or its mutants. Control reactions on the left and right side of the gel were set up 30 minutes before the start of the main reaction and loaded before the first and after the last point of the kinetic experiment. 5 μL reaction aliquots were withdrawn at indicated times and loaded on a running 6% polyacrylamide gel with TBM buffer (89 mM Tris, 89 mM boric acid, 2 mM MgCl_2_). The fraction bound (^32^P-RNA in the binary ^32^P-RNA-Hfq complex or in the ternary ^32^P-RNA-RNA-Hfq complex) was calculated as a proportion of total counts in each lane. To obtain the observed rates of association (*k*_obs_), calculated fractions bound were plotted versus time and fit to single exponential function.

### RNA structure probing

Secondary structure probing experiments were done as previously described [43]. ^32^P-labeled RNAs were denatured prior to the reaction for 1 min at 90°C in water, which was followed by incubation for 10 minutes at room temperature. Then the appropriate reaction buffer used as 5× or 10× was added and mixed with 1 μL of the enzyme. Cleavage reactions were incubated in 10 μL volume in the specific conditions, which are described below. Amounts of the enzymes are indicated on the figures.

RNase T1 reactions in denaturing conditions were carried out for 10 min at 55°C in 7 M urea, 50 mM sodium citrate pH 4.3 buffer, while cleavage in native conditions was performed for 10 min at room temperature in 12 mM Tris–HCl, pH 7.2, 48 mM NaCl, and 1.2 mM MgCl_2_. Reactions with RNase T2 were done at room temperature for 10 minutes in 1× Structure Buffer (10 mM Tris pH 7, 100 mM KCl and 10 mM MgCl_2_; Ambion). Nuclease S1 cleavage was carried out for 15 minutes at room temperature in 1× Reaction Buffer (40 mM sodium acetate pH 4.5, 300 mM NaCl and 2 mM ZnSO_4_; Thermo Scientific), and 0.1 μg of total yeast tRNA (Ambion) was added to RNA prior to reaction.

Each cleavage reaction was quenched by addition of 10 μL buffer STOP (8M urea, 20 mM EDTA). Blank reactions containing 10 μL of 20 nM ^32^P-labeled RNAs were renatured and mixed with 10 μL of the buffer STOP. An alkaline hydrolysis ladder was obtained by incubation of 33 nM final concentration of ^32^P-RNA in formamide for 1 hour at 100°C.

Before loading on the gel, samples were chilled at −20°C. Samples were separated on 10% polyacrylamide gels (19:1 acrylamide/bis-acrylamide ratio) with 8M urea in 1× TBE buffer. The gel was exposed to phosphor screen at −20°C overnight and analyzed using a phosphorimager (Fuji) and Multi Gauge (Fuji) software.

## ACKNOWLEDGMENTS

We thank Sarah Woodson for critical comments on the manuscript. This work was supported by the National Science Centre in Poland (grants No. 2014/15/N/NZ1/03326 to E.M, and No. 2014/15/B/NZ1/03330 to M.O.), KNOW RNA Research Centre in Poznań (No. 01/KNOW2/2014), and Foundation for Polish Science (No. TEAM/2011-8/5) co-financed by the European Union Regional Development Fund within the framework of the Operational Program Innovative Economy.

