## Supplemental data for "Bacterial chaperone protein Hfq facilitates the annealing of sponge RNAs to small regulatory RNAs"

**TABLES**

**Supplemental Table S1. DNA oligonucleotides used in this study.**

| **Name** | **Sequence (5′ → 3′)** |
|---|---|
| AgvB_F | TAATACGACTCACTATAGGCGATAACACAACAATATCAGTATCTCATG |
| AgvB_R | AAAAATGCCCGAATGGGTTCGGGCAATAGCATGAGATACTGATATTGTTG |
| AgvB-U0_R | TGCCCGAATGGGTTCGGGCAATAGCATGAGATACTGATATTGTTG |
| AgvB-U9_R | AAAAAAAAATGCCCGAATGGGTTCGGGCAATAGCATGAGATACTGATATTGTTG |
| GcvB_F | TAATACGACTCACTATAgACTTCCTGAGCCGGAACG |
| GcvB_R | AAAAAAAGCACCGCAATTAGGCGGTGCTACATTAATC |
| GcvB-U1_R | AGCACCGCAATTAGGCGGTGCTACATTAATC |
| 3′ETS$^{leuZ}$_F | TAATACGACTCACTATAgGCCCGGATGGTGGAATCGGT |
| 3′ETS$^{leuZ}$_R | AAAAAAACCACCCGAAGGTGGTTTCACG |
| 3′ETS$^{leuZ}$-U4_R | AAAACCACCCGAAGGTGGTTTCACG |
| RybB_F | TAATACGACTCACTATAGGCCACTGCTTTTCTTTGATGTCCCCATTTTGTGGAGCCCATC |
| RybB_R | AAAAAACCCACCAACCTTGAACCGAAATGGCGGGGTTGATGGGCTCCACAAAATGG |

**Supplemental Table S2. RNA sequences used in this study.**

| Name | Sequence (5′ → 3′) |
|---|---|
| AgvB | ggCGAUAACACAACAAUAUCAGUAUCUCAUGCUAUUGCCCGAA CCCAUUCGGGCAUUUUU |
| AgvB-$U_0$ | ggCGAUAACACAACAAUAUCAGUAUCUCAUGCUAUUGCCCGAA CCCAUUCGGGCA |
| AgvB-$U_9$ | ggCGAUAACACAACAAUAUCAGUAUCUCAUGCUAUUGCCCGAA CCCAUUCGGGCAUUUUUUUUU |
| GcvB | gACUUCCUGAGCCGGAACGAAAAGUUUUAUCGGAAUGCGUGUU CUGGUGAACUUUUGGCUUACGGUUGUGAUGUUGUGUUGUUGU GUUUGCAAUUGGUCUGCGAUUCAGACCAUGGUAGCAAAGCUA CCUUUUUUCACUUCCUGUACAUUUACCCUGUCUGUCCAUAGUG AUUAAUGUAGCACCGCCUAAUUGCGGUGCUUUUUUU |
| GcvB-$U_1$ | gACUUCCUGAGCCGGAACGAAAAGUUUUAUCGGAAUGCGUGUU CUGGUGAACUUUUGGCUUACGGUUGUGAUGUUGUGUUGUUGU GUUUGCAAUUGGUCUGCGAUUCAGACCAUGGUAGCAAAGCUA CCUUUUUUCACUUCCUGUACAUUUACCCUGUCUGUCCAUAGUG AUUAAUGUAGCACCGCCUAAUUGCGGUGCU |
| 3′ETS$^{leuZ}$ | gGCCCGGAUGGUGGAAUCGGUAGACACAAGGGAUUUAAAAUCC CUCGGCGUUCGCGCUGUGCGGGUUCAAGUCCCGCUCCGGGUAC CAUGGGAAAGAUAAGAAUAAAAUCAAAGCAAUAAGCAGUGUC GUGAAACCACCUUCGGGUGGUUUUUUU |
| 3′ETS$^{leuZ}$-$U_4$ | gGCCCGGAUGGUGGAAUCGGUAGACACAAGGGAUUUAAAAUCC |

| | |
|---|---|
| | CUCGGCGUUCGCGCUGUGCGGGUUCAAGUCCCGCUCCGGGUAC CAUGGGAAAGAUAAGAAUAAAAUCAAAGCAAUAAGCAGUGUC GUGAAACCACCUUCGGGUGGUUUU |
| RybB | gGCCACUGCUUUUCUUUGAUGUCCCCAUUUUGUGGAGCCCAUC AACCCCGCCAUUUCGGUUCAAGGUUGAUGGGUUUUUU |

**FIGURE LEGENDS**

**Figure S1. Secondary structure of GcvB sRNA.**

Left: $^{32}$P-labeled GcvB structure probing with RNases indicated above the lanes. T1 D and T1 N denote probing with RNase T1 in denaturing or native conditions, respectively. The numbers on the left indicate positions of guanosine residues. Blank points out untreated control and OH- denotes formamide ladder. Right: secondary structure of GcvB sRNA predicted by RNAstructure software based on the data from structure probing experiments showed on the left. Guanosine residues not cleaved by RNase T1 in native conditions were constrained as double-stranded (green circles). Residues cleaved with RNase T2 or Nuclease S1 were constrained as single-stranded (red circles). Residues involved in base-pairing with AgvB are marked in bold font.

**Figure S2. Hfq-dependent RybB-3′ETS$^{leuZ}$ association.**

The kinetics of $^{32}$P-RybB annealing to 3′ETS$^{leuZ}$ in the presence of 3 nM Hfq. Experiment was performed as in Fig. 3 except that the gel was not run in TBM, but in TBE buffer that allowed migration of RybB-Hfq complexes. Free $^{32}$P-RybB is marked as R, 3′ETS$^{leuZ}$ as L, and Hfq as H. Combinations of those letters refer to their complexes.

**Figure S3. The annealing rates of AgvB-GcvB pair depends on the length of 3′ oligo(U) tail.**

(A-E) The kinetics of $^{32}$P-AgvB annealing to GcvB (and their variants with extended or removed 3′ terminal uridines) in the presence of 9 nM Hfq; (A) AgvB-GcvB, (B) AgvB-$U_9$-GcvB, (C) AgvB-$U_0$-GcvB, (D) AgvB-GcvB-$U_1$, (E) AgvB-$U_9$-GcvB-$U_1$.

(F) The data from A-E were plotted versus time and fitted with single exponential equation. Average $k_{obs}$ values are presented on Fig. 5A.

**Figure S4. The analysis of second order rate kinetics for AgvB-GcvB annealing.**

(A) (left) The fraction $^{32}$P-AgvB annealed to GcvB at indicated Hfq concentrations plotted as a function of time fitted to monophasic rate equation. (right) The average rates of $^{32}$P-AgvB association to GcvB at indicated Hfq concentrations.

(B) (left) The fraction $^{32}$P-AgvB-$U_9$ annealed to GcvB at indicated Hfq concentrations plotted as a function of time fitted to monophasic rate equation. (right) The average rates of $^{32}$P-AgvB-$U_9$ association to GcvB at indicated Hfq concentrations.

### FIGURE S1

A

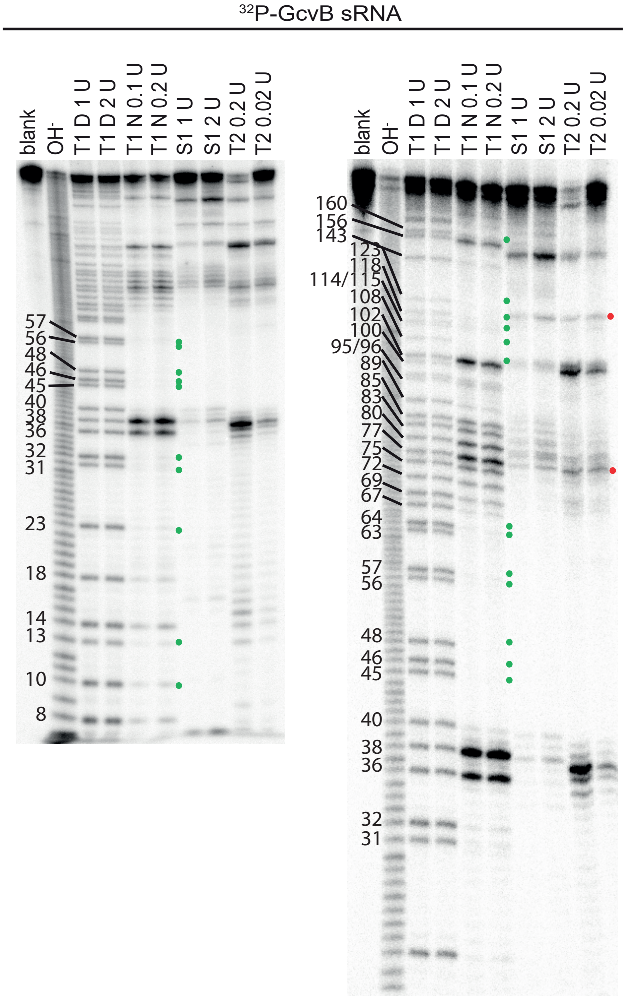

$^{32}$P-GcvB sRNA
blank
OH-
T1 D 1 U
T1 D 2 U
T1 N 0.1 U
T1 N 0.2 U
S1 1 U
S1 2 U
T2 0.2 U
T2 0.02 U
57
56
48
46
45
40
38
36
32
31
23
18
14
13
10
8
160
156
143
123
118
114/115
108
102
100
95/96
89
85
83
80
77
75
72
69
67
64
63
57
56
48
46
45
40
38
36
32
31


B

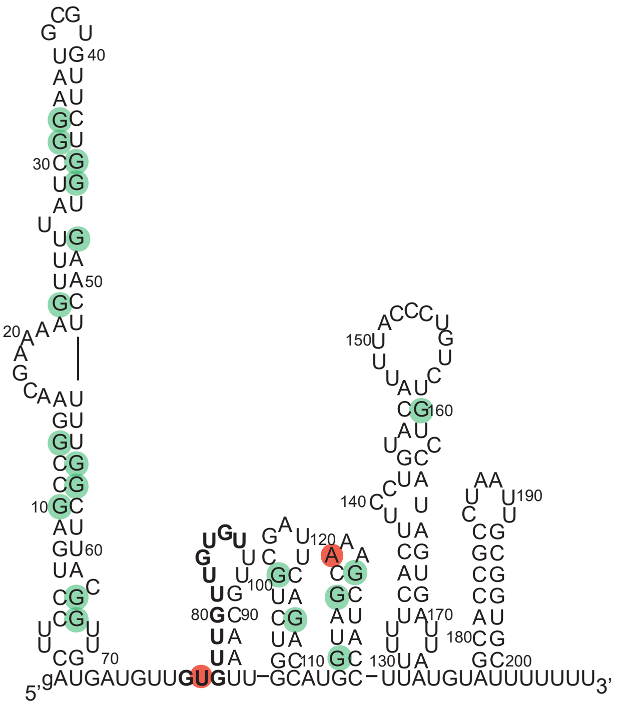

5'
gAUGAUGUUGUGUU–GCAUGC–UUAUGUAUUUUUUU
3'
10
20
30
40
50
60
70
80
90
100
110
120
130
140
150
160
170
180
190
200

**FIGURE S2**

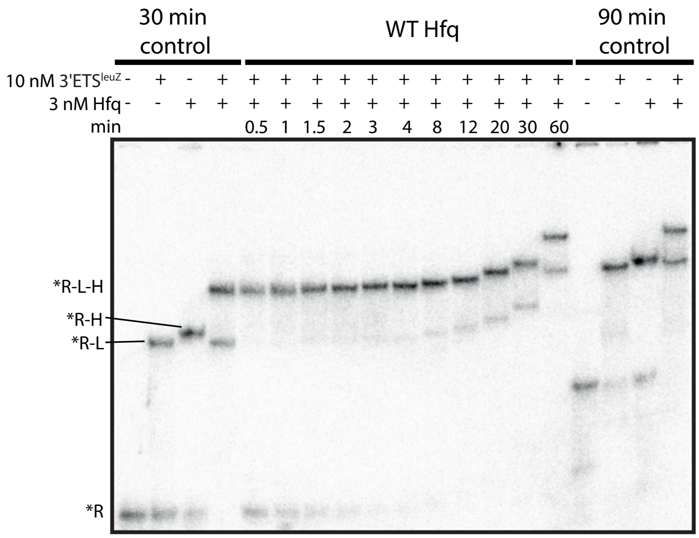

30 min control
WT Hfq
90 min control
10 nM 3'ETS$^{leuZ}$ - + - + + + + + + + + + + + + - + - +
3 nM Hfq - - + + + + + + + + + + + + + - - + +
min 0.5 1 1.5 2 3 4 8 12 20 30 60
*R-L-H
*R-H
*R-L
*R

### FIGURE S3

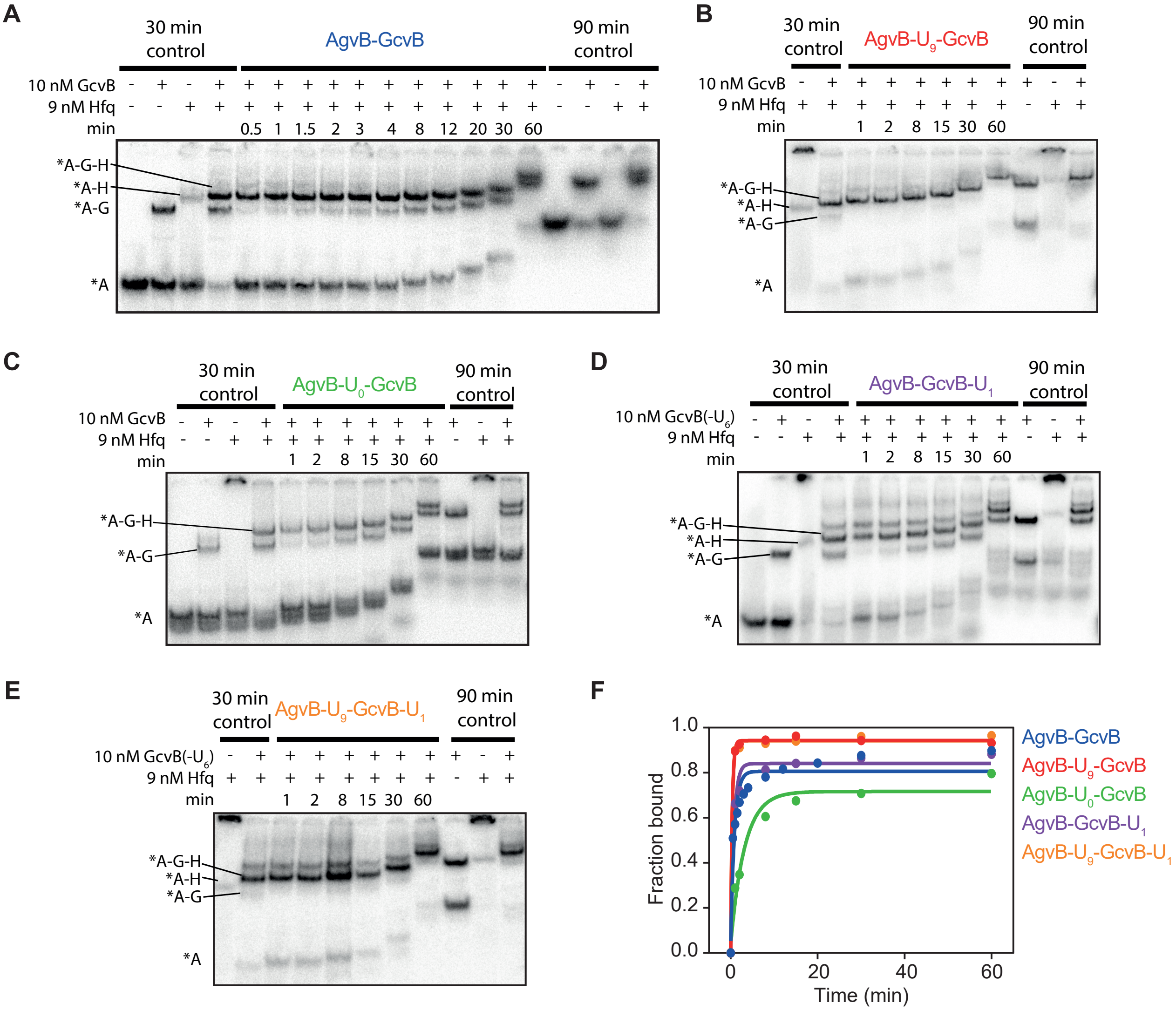

A
30 min control
AgvB-GcvB
90 min control
10 nM GcvB
9 nM Hfq
min
0.5 1 1.5 2 3 4 8 12 20 30 60
*A-G-H
*A-H
*A-G
*A
B
30 min control
AgvB-U9-GcvB
90 min control
10 nM GcvB
9 nM Hfq
min
1 2 8 15 30 60
*A-G-H
*A-H
*A-G
*A
C
30 min control
AgvB-U0-GcvB
90 min control
10 nM GcvB
9 nM Hfq
min
1 2 8 15 30 60
*A-G-H
*A-G
*A
D
30 min control
AgvB-GcvB-U1
90 min control
10 nM GcvB(-U6)
9 nM Hfq
min
1 2 8 15 30 60
*A-G-H
*A-H
*A-G
*A
E
30 min control
AgvB-U9-GcvB-U1
90 min control
10 nM GcvB(-U6)
9 nM Hfq
min
1 2 8 15 30 60
*A-G-H
*A-H
*A-G
*A
F
Fraction bound
Time (min)
AgvB-GcvB
AgvB-U9-GcvB
AgvB-U0-GcvB
AgvB-GcvB-U1
AgvB-U9-GcvB-U1

FIGURE S4

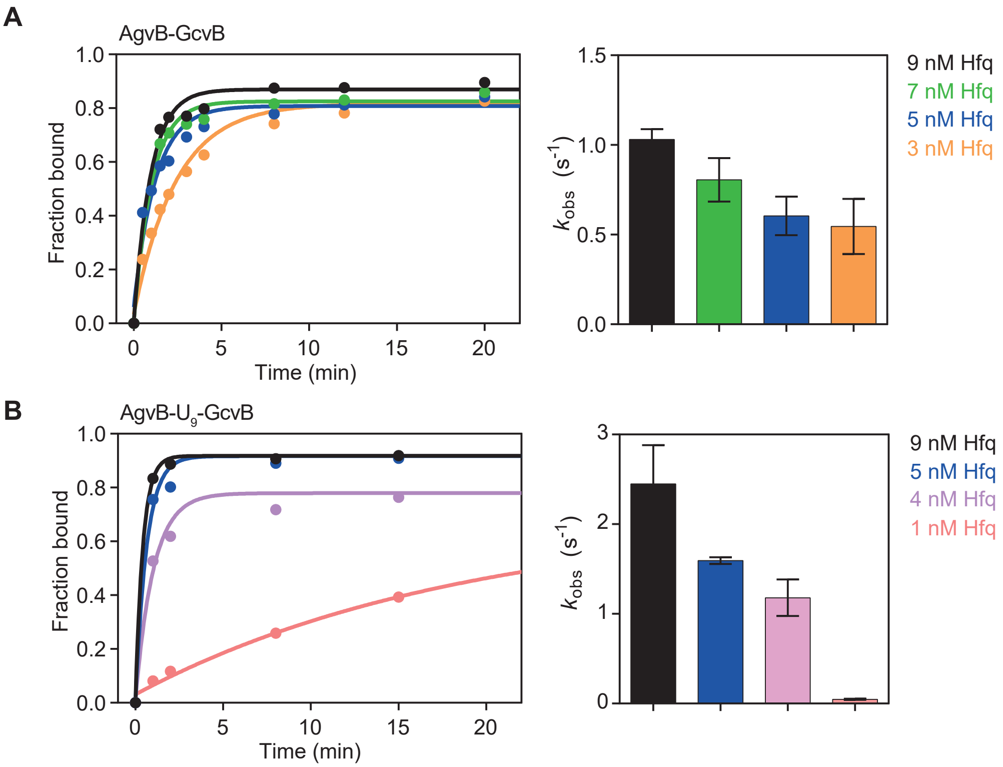
A
AgvB-GcvB
Fraction bound
Time (min)
$k_{obs}$ ($s^{-1}$)
9 nM Hfq
7 nM Hfq
5 nM Hfq
3 nM Hfq
B
AgvB-$U_9$-GcvB
Fraction bound
Time (min)
$k_{obs}$ ($s^{-1}$)
9 nM Hfq
5 nM Hfq
4 nM Hfq
1 nM Hfq
